# A single meiotic product dominates inheritance in *Mucor circinelloides*

**DOI:** 10.64898/2026.10.23.761428

**Authors:** Carlos Pérez-Arques, María Isabel Navarro-Mendoza, Ziyan Xu, Lysett Wagner, Grit Walther, Joseph Heitman

**Affiliations:** Department of Molecular Genetics and Microbiology, Duke University School of Medicine, NC 27710, USA; Department of Genetics and Microbiology, Faculty of Biology, University of Murcia, 30100 Murcia, Spain; National Reference Center for Invasive Fungal Infections, Leibniz Institute for Natural Product Research and Infection Biology, Hans Knöll Institute, 07745 Jena, Germany

## Abstract

Meiosis is a major source of genetic diversity, yet it remains poorly understood in many pathogenic fungi. In Mucorales, the causative agents of mucormycosis, sexual reproduction has been difficult to study, and reported inheritance patterns have appeared inconsistent with conventional meiosis. Here, meiosis in *Mucor circinelloides* was demonstrated at genome-wide resolution by generating telomere-to-telomere chromosome assemblies from compatible mating types and analyzing parental markers distributed across the genome. The results reveal meiotic recombination and independent chromosome assortment. However, the outcome of meiosis is strikingly unconventional: all sexual spores derived from a single zygospore share an identical nuclear genotype, even when germination produces distinct sporangia. This pattern suggests that the viable progeny of each zygospore is derived from a single meiotic product, rather than from the four products expected under conventional meiosis. Mitochondrial inheritance is also uniparental, with progeny mitochondria inherited from the (+) mating-type parent. Together, these findings show that sexual reproduction in *M. circinelloides* entails meiotic recombination, but restricts inheritance to a single viable nuclear product and a single mitochondrial lineage. This decoupling of recombination from equal meiotic contribution reveals a hidden layer of inheritance control in Mucorales, showing how pathogenic fungi can reshuffle chromosomes while producing genetically uniform sibling progeny, an outcome reminiscent of female meiosis in animals.

## Introduction

Sexual reproduction is a key hallmark of eukaryotic life (Speijer et al., 2015). At its core, meiosis and recombination reshape genomes, generating diversity, purging deleterious mutations, and driving rapid adaptation (McDonald et al., 2016). Meiosis achieves this during its first division – meiosis I– through the independent assortment of homologous chromosomes and crossing over. After sister chromatid separation in the second division, meiosis II forms four haploid meiotic products, each carrying a unique combination of parental alleles. Despite its importance, sexual reproduction and meiosis remain understudied in many eukaryotic microbes (Gaunt et al., 2003; Josling et al., 2018; Yadav et al., 2023; Yildirir et al., 2020). For instance, the recently characterized sexual cycle of the human pathogen *Aspergillus fumigatus* (O’Gorman et al., 2009) revealed an unusually high crossover rate that may generate novel haplotypes contributing to drug resistance and pathogenesis (Auxier et al., 2023). Other fungal pathogens such as *Candida albicans* lack a conventional sexual cycle and instead, undergo a parasexual cycle with systematic loss of duplicated chromosomes in a parameiotic process (Bennett, 2003; Pontecorvo et al., 1953). In many pathogens multiple sexual strategies may coexist, including unisexual and pseudosexual reproduction of *Cryptococcus neoformans* leading to uniparental nuclear inheritance (Lin et al., 2005; Yadav et al., 2021). As fungal infections gain recognition by the World Health Organization as a major global health threat (Fisher & Denning, 2023), understanding the mechanisms that promote genomic diversity may reveal how different sexual strategies contribute to host adaptation and the evolution of drug resistance.

Among these emerging threats, mucormycoses stand out as rare yet deadly invasive fungal infections caused by molds of the order Mucorales (Skiada et al., 2025). These fungi are notorious for rapid host angioinvasion and unusually high drug resistance to common antifungal therapies. As a result of treatment failure, mucormycoses can escalate to disseminated infections with mortality rates approaching 90% (Cornely et al., 2019). Despite their clinical significance, little is known about sexual reproduction and meiosis in pathogenic Mucoralean species, partly because completing the sexual cycle under laboratory conditions is exceptionally challenging (Pérez-Arques et al., 2025; Wagner et al., 2020). Curiously, most of what we understand about Mucoralean sexual reproduction is focused on studies of non-pathogenic species rather than the pathogenic ones.

Studies in these model organisms have revealed fundamental features of fungal sexual reproduction, including the first observation of mating and sexual spore formation (Ehrenberg, 1820), heterothallism (Blakeslee, 1904), and the role of diffusible pheromones (Burgeff, 1924). In the Mucorales, sexual identity is determined by a high-mobility group transcription factor encoded by two idiomorphic alleles, *sexP* and *sexM*, corresponding to the (+) and (–) mating types, respectively (Idnurm et al., 2008). Hyphae of opposite mating types sense each other via pheromone signaling and then form specialized hyphal branches called zygophores. Upon contact, zygophores differentiate into gametangia, separated from the mycelium by septa, and fuse to form the zygospore (Blakeslee, 1904). From the thousands of nuclei present within the zygospore, only two undergo karyogamy to form a single diploid nucleus, which then proceeds through meiosis to generate four genetically distinct haploid products after a dormancy period (Cerdá-Olmedo, 1975; Chaudhary et al., 2013; Eslava et al., 1975). As the zygospore germinates, it forms a germsporangium that releases haploid germsporangiospores, the sexual progeny.

The existence of canonical meiosis in the Mucorales has long been debated. Early observations in *Phycomyces blakesleeanus* revealed sex heterokaryons and inheritance patterns that defied Mendelian ratios (Cerdá-Olmedo, 1975; Mehta & Cerdá-Olmedo, 2001), challenging whether conventional meiosis occurs. This uncertainty was partially resolved when a genetic map confirmed genome-wide meiotic recombination in *P. blakesleeanus* (Chaudhary et al., 2013). Nonetheless, unusual inheritance patterns persist in *Mucor* species, where all progeny from a single zygospore consistently share the same mating type (Gauger, 1965; Köhler, 1935; Sjöwall, 1945) and the same set of six genetic markers (James & Gauger, 1982). These anomalies suggest alternatives to conventional sexual reproduction and meiosis, raising fundamental questions about how Mucorales generate genetic diversity.

Here in this study, we combined recent sequencing technologies and classical genetics to rediscover the long-neglected sexual cycle of the human pathogen *Mucor circinelloides*. Our results reveal genome-wide meiotic recombination yet remarkably, only one meiotic product survives meiosis to define the genotype of all progeny. These findings challenge classical expectations of balanced meiotic contribution and reshape our understanding of genetic diversity in these pathogenic fungi with striking parallels to female meiosis in animals.

## Results

Accurate genome assemblies are essential for resolving chromosome-scale recombination patterns and understanding the genomic architecture that shapes meiosis. Previous draft assemblies of *Mucor circinelloides* were incomplete and lacked chromosome-level resolution, which complicated the analysis of genetic variation and meiotic recombination. This limitation was addressed by generating complete, telomere-to-telomere genome assemblies and annotations from the (+) and (–) mating types of *M. circinelloides* (Table S1), combining Oxford Nanopore Technologies (ONT) long reads and proximity ligation techniques to capture chromatin conformation (Hi-C). To characterize chromosome structure, we identified centromeres by mapping previously generated histone H3 lysine 9 dimethylation (H3K9me2) profiles and correlating these data with the presence of a centromere-specific DNA motif next to a highly AT-rich stretch, found within large heterochromatic regions devoid of protein-coding genes and enriched with clusters of centromeric retrotransposable elements termed Grem-LINE1s (Figure 1a, zoomed in Figure 1b). Each assembly consists of ten complete chromosomes ending in canonical telomeric repeats (5’-TTAGGG-3’) and the majority display an acrocentric architecture (Table S1). We also identified three chromosomes carrying ribosomal DNA (rDNA) arrays located near the centromere. Importantly for downstream applications, both mating-type genomes are largely syntenic (Figure 1a, alignments), ensuring mating compatibility and facilitating subsequent genetic variation analyses.

**Figure 1.**
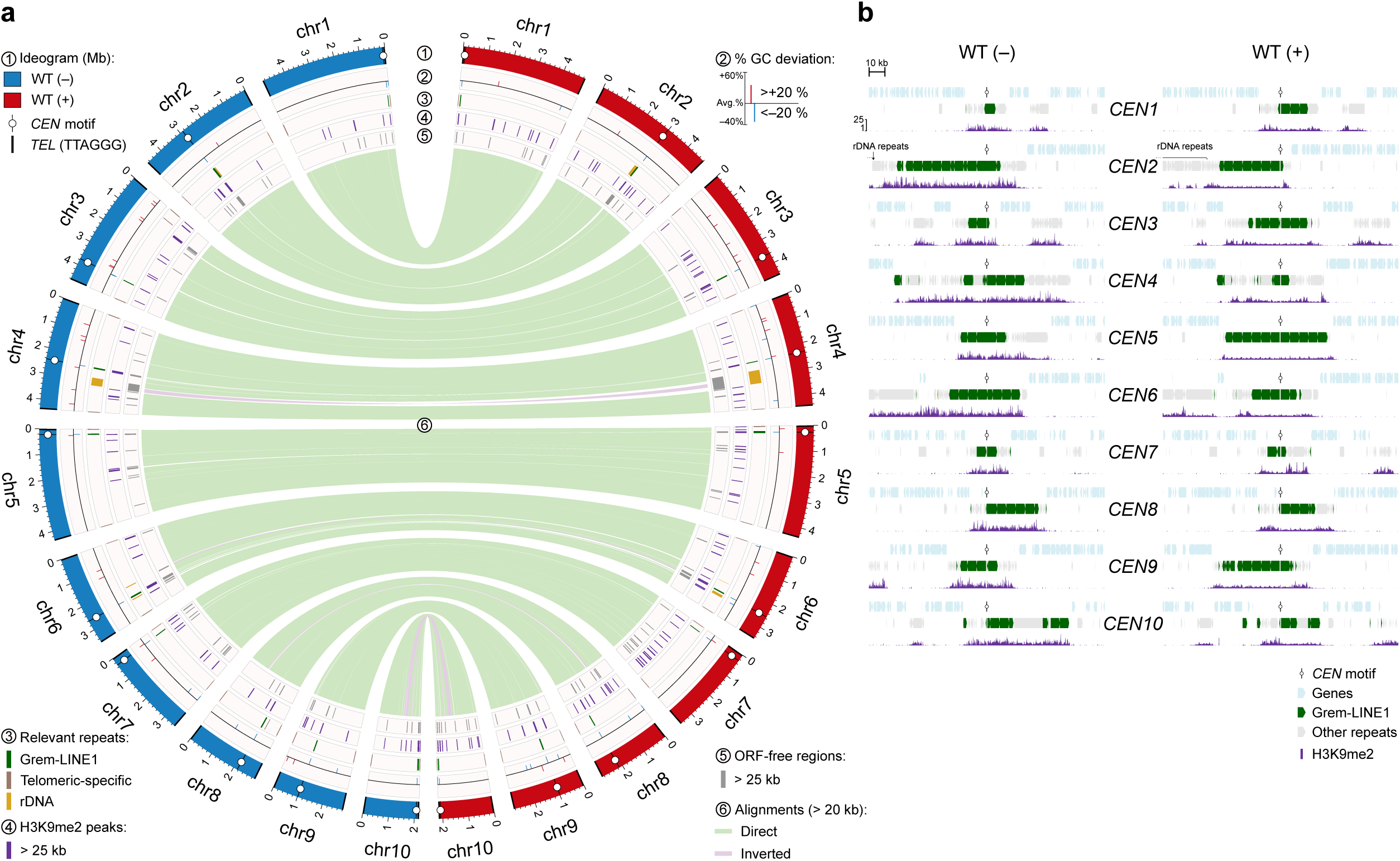
*Mucor circinelloides* PS15 opposite mating-type genomes exhibit high synteny. **(a)** A circular plot depicts the complete, telomere-to-telomere genome assemblies of *M. circinelloides* (–) and (+) mating types, with annotated features defined by the legend around the plot. Each mating-type ideogram is color-coded and displays centromeric-and telomeric-specific (TTAGGG) sequence motifs. GC content is represented as the percentage deviation (>±20%) from the genome-wide average. Location-specific repeated elements (Grem-LINE1 enriched in centromeres, telomere-specific elements, and rDNA clusters), as well as large regions (>25 kb) enriched in H3K9me2 and devoid of open reading frames (ORFs) are shown color-coded. Alignment links highlight the high synteny and contiguity between the two assemblies. **(b)** Genomic plots of pericentric regions (150 kb, oriented 5′→3′ on the top strand) show the presence of a conserved *CEN* motif, arrays of Grem-LINE1 and other repeats, and large heterochromatic, ORF-free domains, indicating the location of ten centromeres per genome assembly.

Sexual reproduction in the Mucorales begins when sexually determined, haploid zygophores from the (+) and (–) mating types come into contact and develop into gametangia. After plasmogamy, these structures mature into zygospores containing hundreds of nuclei from both mating types. Two opposite-mating-type nuclei fuse to form a diploid nucleus that then undergoes meiosis, producing haploid meiotic products (Figure 2a). Dormancy ends with zygospore germination, with each one forming a germsporangium that contains germsporangiospores (Figure 2b). To assess whether the genetic diversity of germsporangiospores (for simplicity hereafter referred to as spores) from a single zygospore reflects contributions from multiple meiotic products, we mated two opposite-mating-type wildtype *M. circinelloides* <u>p</u>hylogenetic <u>s</u>pecies <u>15</u> (PS15) strains under laboratory conditions (Figure 2a-b).

**Figure 2.**
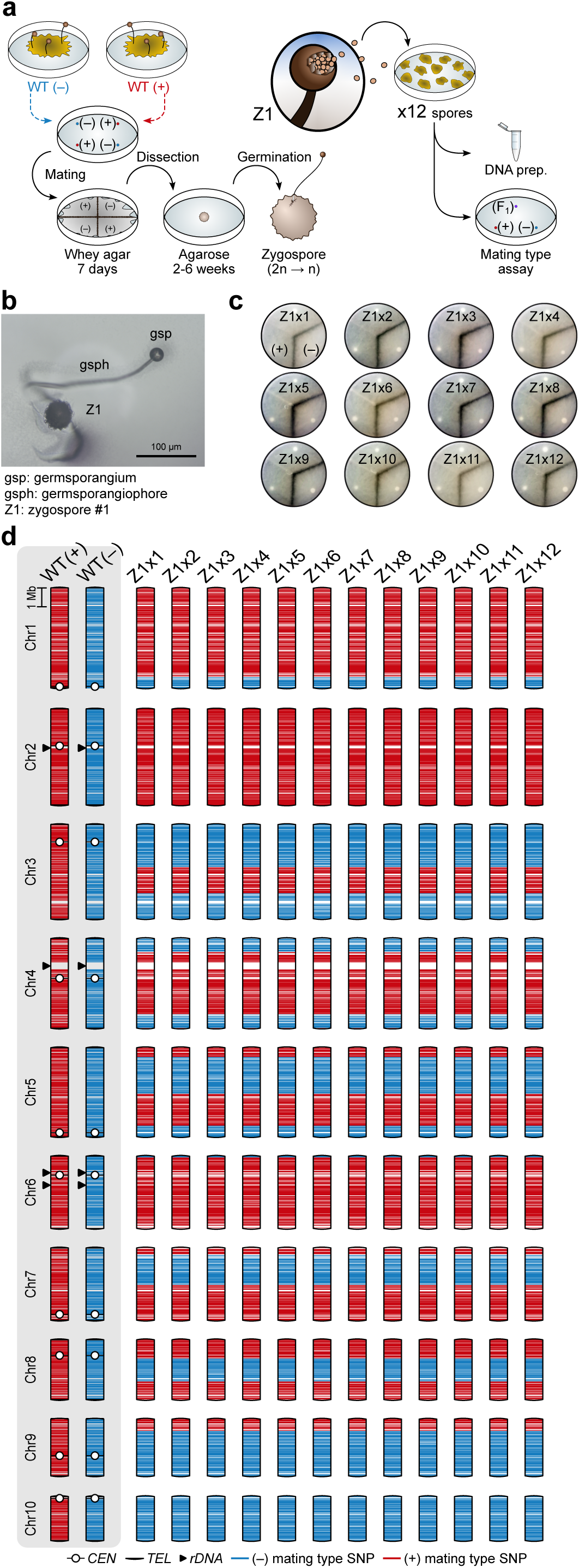
Germsporangiospores derived from a single zygospore exhibit identical meiotic recombination patterns. **(a)** Overview of the experimental procedure for isolating F1 progeny. Briefly, fresh spores of opposite mating types were inoculated equidistantly to ensure compatible mating. After ∼1 week of incubation in the dark, mature zygospores appearing as a dark-pigmented strip at mating-compatible intersections were manually dissected and transferred to individual plates until germination (dormancy stage of 2 to 6 weeks). Upon zygospore germination, germsporangiospores were collected and spread to isolate single-spore colonies. Twelve colonies were selected for mating-type determination and DNA preparations. **(b)** Composite light microcospy images (z-stack, z=3) showing the typical morphology of a germinating zygospore. **(c)** Mating type assay plates demonstrate that all twelve single-spore colonies (Z1×1 to Z1×12) derived from the same zygospore (Z1) share the same mating type (+). Each colony was confronted with both (+) and (–) wild-type strains (WT). Dark-pigmented mycelial strips at colony intersections with the WT (–) indicate successful mating and enrichment of zygospores. **(d)** Schematic chromosome representations indicate SNP inheritance patterns in the progeny, with + mating type alleles in red and – mating type alleles in blue. Every progeny from the same zygospore (Z1) displays identical patterns of independent chromosome assortment and meiotic recombination. Parental WT (+) and (–) chromosomes show centromeres, telomeres, and rDNA repeats for reference.

We isolated spores from a single zygospore (Z1), selecting twelve for subsequent analyses. Backcrossing these progeny revealed that every spore analyzed harbored the same mating-type [Figure 2c, (+) mating type)], suggesting that spores derived from the same zygospore may be genetically identical. To test this, we performed Illumina whole-genome sequencing on mycelium from individual spores and identified 18,587 single nucleotide polymorphisms (SNPs) distinguishing the two parental genomes. These SNPs served as genetic markers that demonstrated that all twelve spores from the same zygospore were genetically identical, resolving meiotic recombination and chromosome assortment patterns across the progeny (Figure 2d). Analysis of a second zygospore (Z3) revealed that all five spores examined were also genetically identical, generalizing these findings.

During zygospore isolation, a small proportion of germinating zygospores exhibited sympodial branching germsporangiophores (Figure 3a, b). We hypothesized that distinct meiotic products could drive separate branches, such that spores from separate germsporangia might carry different genotypes. To test this, 12 spores from each of two distant germsporangia of the same zygospore (Z2.g1 and Z2.g2) were dissected. Whole-genome data revealed identical meiotic recombination and chromosome assortment patterns across both germsporangia (Figure 3c). These findings were replicated in another branching zygospore, indicating that spores from the same zygospore are genetically identical, regardless of germsporangiophore branching.

**Figure 3.**
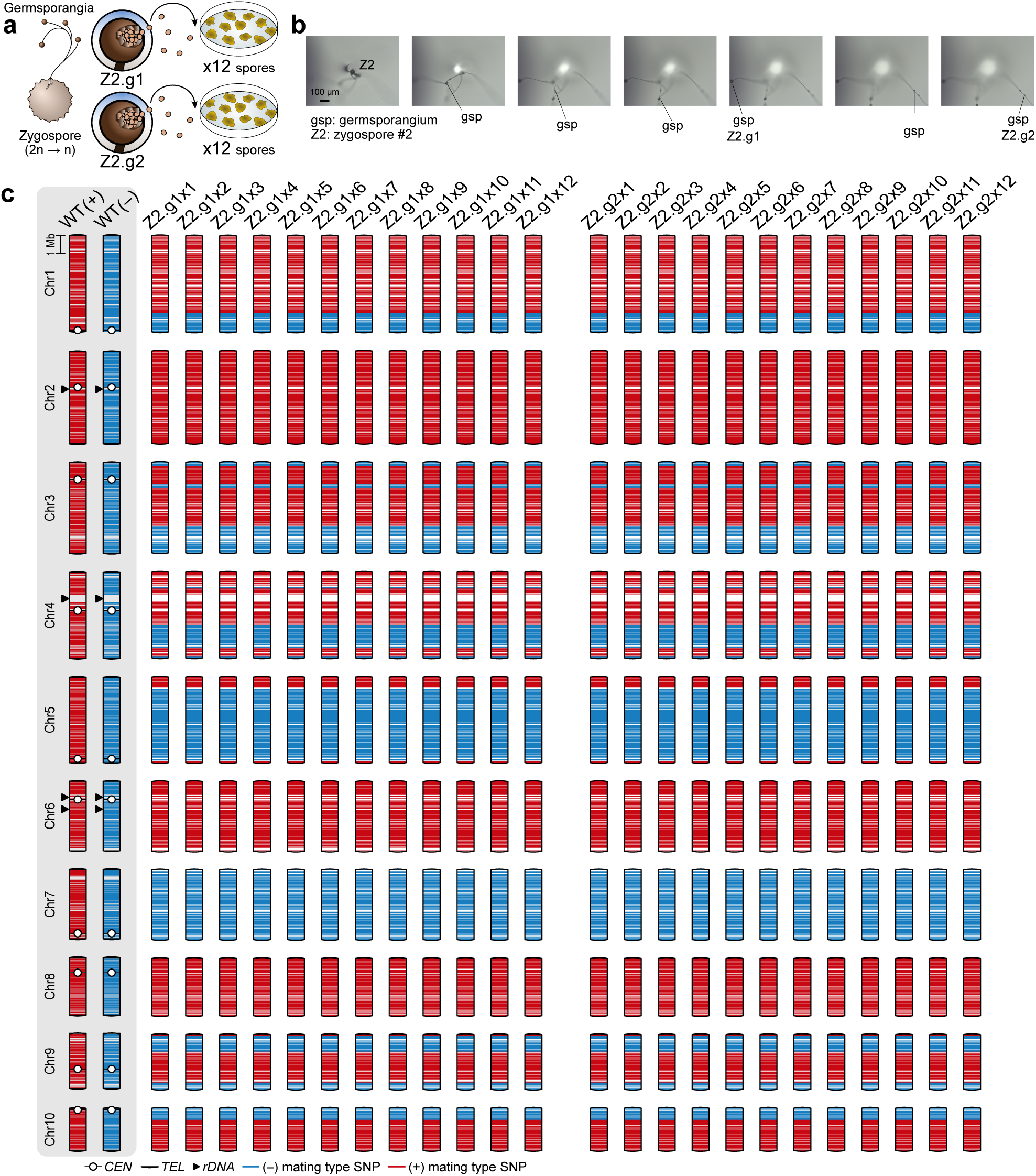
Germsporangiospores from branching germsporangia are genetically identical. **(a)** Illustration of a branching germinating zygospore (Z2) and the workflow to isolate F1 progeny from separate germsporangia (g1 and g2). **(b)** Z-stack light microscopy images of a branching germsporangiophore arising from a single zygospore (Z2). The two most distal germsporangia (Z2.g1 and Z2.g2) were collected separately and analyzed. **(c)** Distribution of paternal SNPs across twelve single-spore samples from the two distinct germsporangia (Z2.g1×1–12 and Z2.g2×1–12) stemming from the same zygospore shows identical patterns of chromosome assortment and meiotic recombination.

Assuming four genetically distinct meiotic products contribute equally to spore genotype, the probability of *n* spores being genetically identical is given by *P* = 4 × (1⁄4)*^n^* = (1⁄4)*^n^*^−1^. Under this model, testing with our most conservative sampling (n = 5) yields a probability of 0.391% (Table 1). The combined probability of observing genetically identical spores across all four zygospores is 5.05 x 10^-29^, strongly supporting that progeny from the same zygospore derive from a single meiotic product.

**Table 1.** Probability (*P*) that spores from each zygospore exhibit identical genotypes under the null hypothesis that all four meiotic products contribute equally. Combined probability assumes the four zygospores represent independent trials.

|  | **Z1 (n = 12)** | **Z2 (n = 24)** | **Z3 (n = 5)** | **Z4 (n = 10)** | **Combined** |
| --- | --- | --- | --- | --- | --- |
| ***P*** | $2.38 \times 10^{-7}$ | $1.42 \times 10^{-14}$ | $3.91 \times 10^{-3}$ | $3.81 \times 10^{-6}$ | $5.05 \times 10^{-29}$ |

To expand our analysis, we examined progeny from 52 additional zygospores collected in an earlier study. From each zygospore, hundreds of spores were pooled and sequenced to evaluate genotype uniformity, reasoning that the presence of heterozygosity at informative SNP markers would suggest mixed progeny genotypes deriving from more than one meiotic product. Instead, bulk sequencing results consistently revealed high levels of homozygosity (≥ 95%) across all SNP positions that define the meiotic recombination pattern. We applied a binomial probability model under the null hypothesis that spores were randomly sampled from four genetically distinct meiotic products, based on the conservative assumption that each spore pool contained only two spores [*P* = 4 × (1⁄4)^2^ = 1⁄4]. Each zygospore was treated as an independent trial, and a pool was considered to originate from a single meiotic product if it exhibited > 95% homozygosity. From this model, the probability of observing such uniformity across all 52 zygospores is *P* = 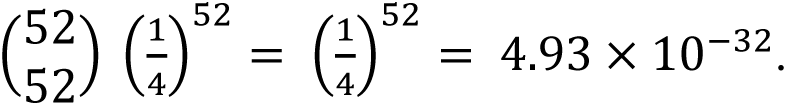

Our analyses show that meiosis in a single zygospore results in spores with genetically identical nuclear genomes. However, additional sources of genetic variation could influence progeny genotypes. Mitochondrial genomes, inherited independently of nuclear chromosomes, represent a source of such variation. The type of mitochondrial inheritance in Mucorales has only been characterized in *Phycomyces blakesleeanus*, which exhibits mitochondrial uniparental inheritance (UPI). Notably, all four meiotic products survive and contribute to progeny genotype in *P. blakesleeanus*, suggesting that mitochondrial inheritance dynamics could also differ in *M. circinelloides*. To investigate this, we assembled and annotated the mitochondrial genomes of both mating types (Figure 4a), identifying 20 SNPs to track mitochondrial inheritance and assess its contribution to spore genotypes. Mitochondrial genomes were unambiguously inherited from the (+) mating-type parent across all progeny analyzed in this study (Figure 4b), including spore pool samples from previously collected zygospores (Figure 4c). To provide a rapid, reliable, and cost-efficient framework for future mtDNA analyses, we designed a restriction fragment length polymorphism (RFLP) approach (Figure 4a). This RFLP analysis confirmed mitochondrial UPI from the (+) mating-type parent in all of the 47 progeny from *M. circinelloides* examined. Together, these results demonstrate that a single meiotic product dominates inheritance in *M. circinelloides*, yielding genetically uniform progeny across both nuclear and mitochondrial genomes. This pattern contrasts with classical expectations of balanced meiotic contribution and suggests that mechanisms of inheritance in the pathogenic *Mucor circinelloides* species complex may restrict genetic diversity in ways that could profoundly influence their evolution and adaptation.

**Figure 4.**
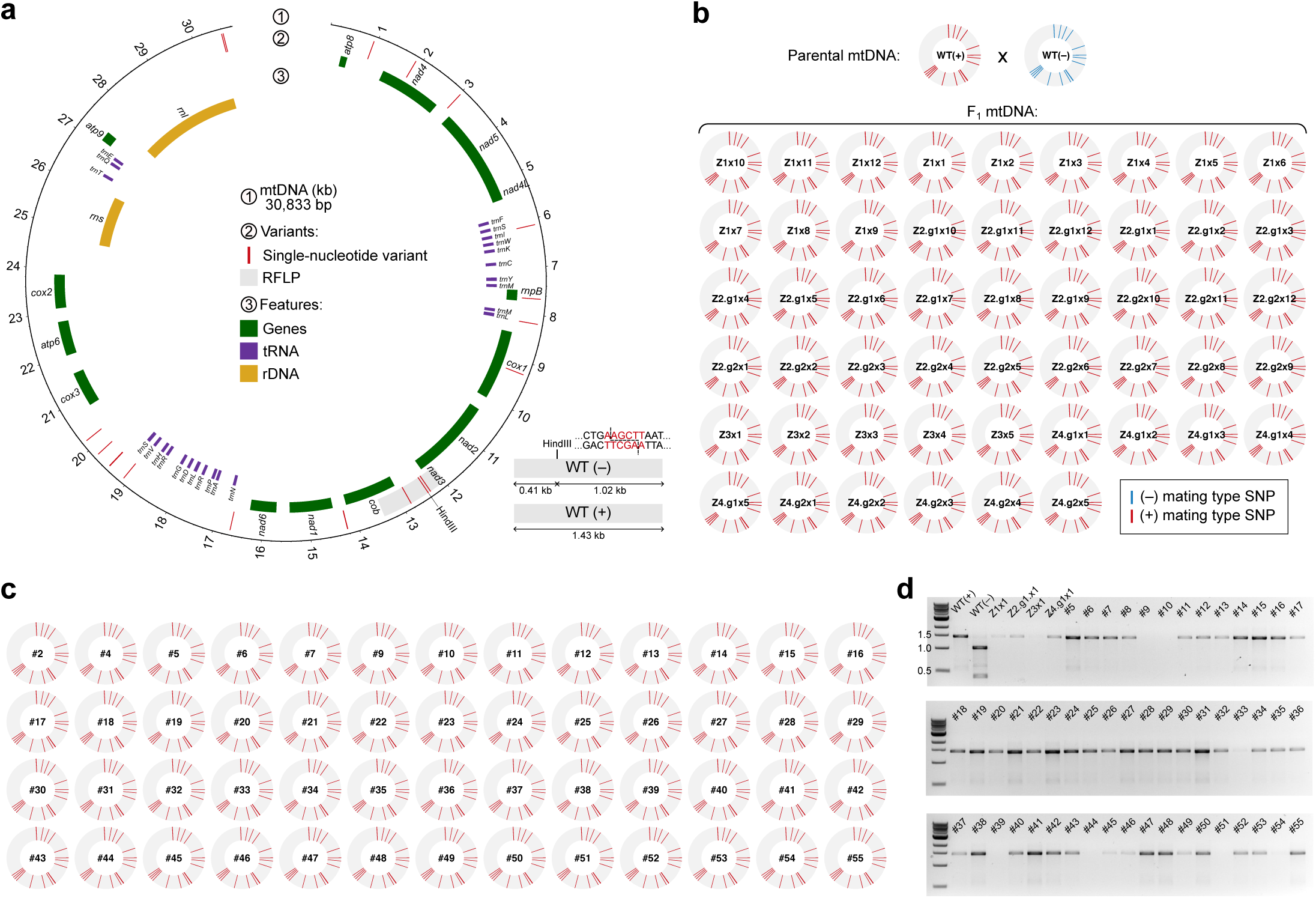
*M. circinelloides* displays mitochondrial uniparental inheritance from the (+) mating-type parent. **(a)** Circular map representing *M. circinelloides* PS15 annotated mitochondrial DNA (mtDNA). The plot highlights identified SNPs between (+) and (–) mtDNA, a HindIII restriction fragment length polymorphism (RFLP), and mtDNA genes, tRNAs, and rRNA genes as defined by the color-coded legend. The expected fragment sizes and HindIII cut site for the RFLP are shown in the lower right corner. **(b, c)** Parental SNP distribution in the progeny mtDNA, with + mating type alleles in red and – mating type alleles in blue. Panel **(b)** shows data from 52 progeny generated by this study; and panel **(c)** shows data from 52 progeny analyzed in a previously published dataset. All 104 progeny mtDNA consistently inherited (+) mating-type alleles across the 20 identified mtDNA SNPs, supporting uniparental mtDNA inheritance from the (+) mating-type parent. **(d)** HindIII RFLP agarose electrophoresis of mtDNA from a representative set of progeny validates the RFLP assay as a rapid screening tool for determining mtDNA inheritance. Parental WT (+) and (–) RFLP patterns and a DNA ladder are included as references.

**Figure 5.**
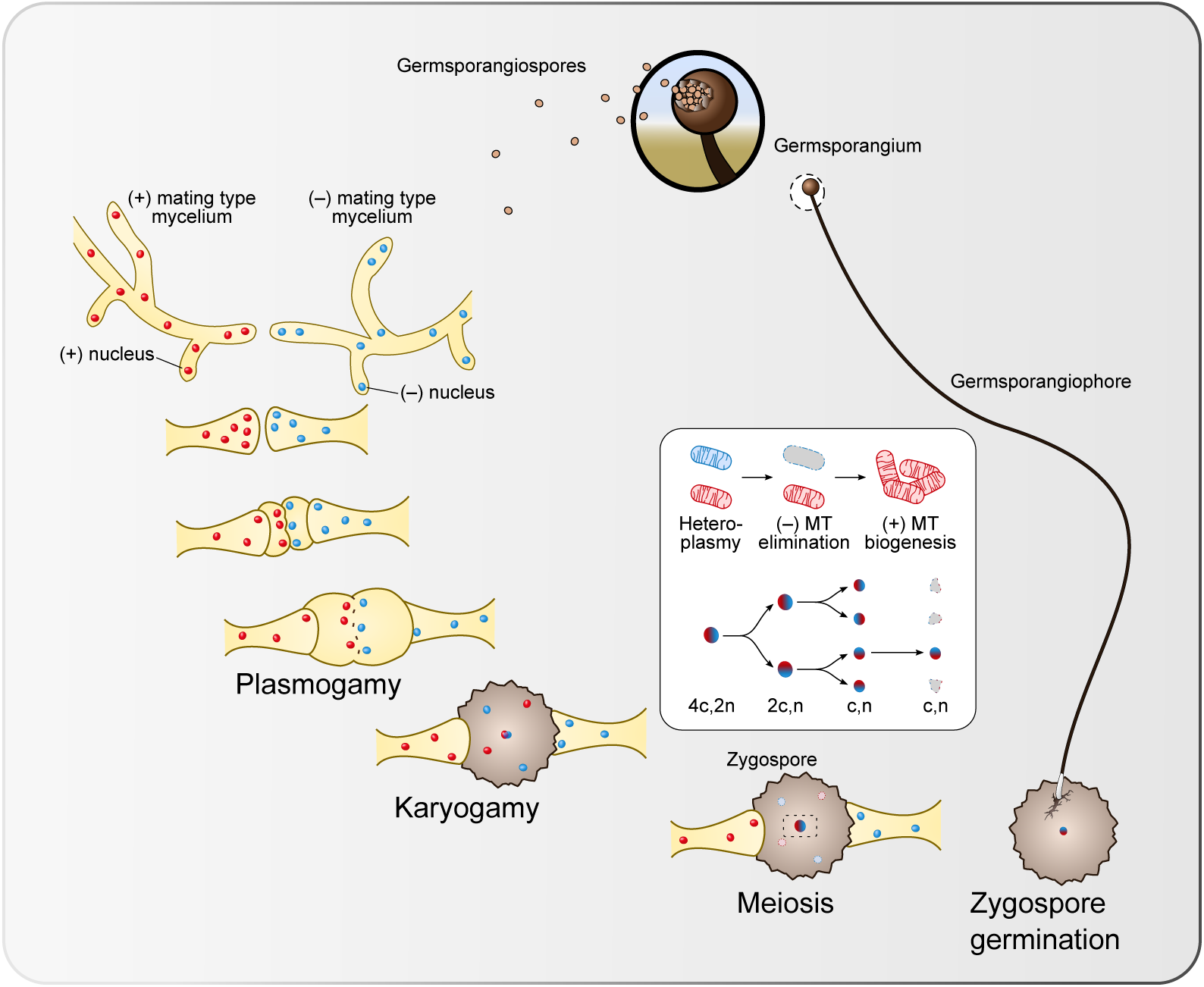
Model of sexual reproduction and genetic inheritance in *M. circinelloides*. Model illustrating the major stages of the *M. circinelloides* sexual cycle. Two opposite-mating-type, non-septate hyphae called zygophores grow toward each other, forming engorged progametangia that fuse to produce gametangia. The gametangia then become separated from the parent hyphae by septa. Plasmogamy follows, generating a prozygosporangium that matures into a zygospore. Within the zygospore, karyogamy occurs between two haploid, opposite-mating-type nuclei, followed by meiosis. Our model hypothesizes that, early in this process, the initial mitochondrial heteroplasmy is resolved through targeted degradation or mitophagy of (–) mating-type mitochondria, accompanied by uniparental mitochondrial inheritance of the (+) mating type. Genetic progeny analysis reveals that only one meiotic product survives, while the other three are lost during development, similar to female meiosis in animals. After a dormancy period, the zygospore germinates into a germsporangiophore that branches into a single or several germsporangia containing genetically identical, haploid germsporangiospores.

## Discussion

Our studies resolve the long-standing mystery of the sexual cycle of the human fungal pathogen *M. circinelloides* by demonstrating that the process includes meiotic recombination and independent chromosomal assortment. Strikingly, our studies reveal that only a single meiotic product survives from each meiotic event. The biological reason for this may be selective forces in nature in which the organism has benefited by reducing inbreeding potential and promoting outcrossing. Given that there are thousands of spores produced within each germsporangium, if all four meiotic products survived there would be ample opportunity for sibling F1 progeny to mate with each other to result in inbreeding. A single genotype surviving each meiotic event ensures that any subsequent and compatible encounter in nature will be with a genetically divergent mating partner. Given that the germsporangium contains hundreds to thousands of identical nuclei, the sole surviving meiotic product nucleus must undergo many rounds of subsequent mitotic division to produce a multitude of spores. This is conceptually similar to post-meiotic rounds of mitosis that occur in other fungi, such as *Neurospora crassa* (which produces octads with eight spores) and *Cryptococcus neoformans*, which can produce hundreds of spore products descended from only four initial meiotic nuclei. There are other examples in the fungal kingdom in which meiosis results in fewer than the expected four meiotic products. One example is the model budding yeast *Saccharomyces cerevisiae*. While this organism typically produces tetrads containing four meiotic products, a range of outcomes is observed with some wild yeast strains in which triads (three spores) and dyads (two spores) are also produced (Yang et al., 2026). Moreover, when acetate is omitted from standard yeast sporulation media, or sporulation is interrupted by transfer from media to water, sporulation results in the production of dyads and not tetrads (Davidow et al., 1980; Srivastava et al., 1981; Taxis et al., 2005; Neiman, 2011). These dyads are of a specific genetic composition known as non-sister dyads, and this maximizes the genetic diversity that is possible with only two meiotic products. Under these altered nutritional conditions, two of the four meiotic nuclei are degraded during spore formation, resulting in only two viable meiotic products. Another example in which sporulation results in dyads is in the pathogenic yeast *Candida lusitaniae*. This species is a rare cause of human infections, and while it is a member of the CTG-clade of yeasts like *Candida albicans*, in contrast to *C. albicans* which is diploid *C. lusitaniae* is haploid. Mating between cells of the two opposite mating types results in a diploid zygote that forms an ascus containing two clavate or club-shaped spores. Genetic analysis has shown that the dyads produced by *C. lusitaniae* are non-sister dyads (Reedy et al., 2009). Whether the sporulation media utilized to study *C. lusitaniae* are missing a nutrient, similar to lack of acetate leading to dyads in *S. cerevisiae*, is unknown. One final unusual example is *Lodderomyces elongisporus*, a recently recognized pathogenic yeast that is related to the more common pathogens *Candida parapsilosis*, *Candida orthopsilosis*, and *Candida metapsilosis* (Lockhart et al., 2008). Curiously, *L. elongisporus* is thought to have an extant sexual cycle but produces asci that contain only one spore, which are called monads. Studies remain to be conducted to prove that these are the products of homothallic sexual reproduction and meiosis, but if validated, this would be another fungal example of a species that undergoes sexual reproduction in which only one meiotic product survives from each meiotic event.

The production of only a single meiotic product by Mucorales sexual reproduction is also reminiscent to female meiosis in animals, but with important differences. During gamete development in animals, male meiosis/spermatogenesis result in four viable meiotic products, whereas female meiosis produces only one final viable meiotic product, the mature ovum (Fragouli et al., 2011; Evans & Robinson, 2011). The process of female meiosis is complex and involves different steps at different developmental stages. In humans, the primary oocyte is formed in the fetus during gestation, and this cell enters prophase I and becomes quiescent. During puberty, the primary oocyte undergoes meiosis I, yielding a secondary oocyte that arrests in metaphase II and a second product that is called the first polar body. Subsequently, when the secondary oocyte fuses with a sperm, an influx of calcium triggers completion of meiosis II to form a mature ovum and a second polar body. The first polar body may also undergo cell division to produce a total of three polar bodies in some cases. The polar bodies that are produced as products of female meiosis all degenerate and die. Thus, through this process only one mature female meiotic product survives, which is the ovum. By producing only one meiotic product, oogenesis enables production of a large ovum and two or three much smaller polar bodies. This asymmetric division preserves cytoplasm and organelles in the female gamete while excluding the additional meiotic products. There is no evidence yet for an equivalent cytological mechanism in Mucorales, but given that the products of meiosis in fungi are equivalent in size, we conjecture that production of only one or two meiotic products in fungi is to 1) maximize the chances of producing viable spores when nutrients are limiting, and 2) reducing inbreeding to promote outbreeding. Whether other examples analogous to female meiosis occur in fungi remains to be explored in future studies.

Mitochondrial inheritance adds a second layer to the biased transmission observed during sexual reproduction *in M. circinelloides*. Across all progeny analyzed, mtDNA was inherited uniparentally from the (+) mating-type parent, indicating that sexual development transmits not only a single nuclear meiotic product, but also a single mitochondrial lineage. This pattern mirrors previous work in *P. blakesleeanus*, where mitochondrial inheritance is also directed by mating-type identity and proceeds from the (+) parent, with loss of mtDNA from the (–) parent (Shakya & Idnurm, 2014). The agreement between *M. circinelloides* and *P. blakesleeanus* suggests that (+)-biased mitochondrial inheritance may be a conserved feature of sexual reproduction in Mucorales. Similar cases of mitochondrial UPI or strongly biased mitochondrial inheritance have been described across fungi. In basidiomycetes such as *Ustilago maydis* and *C. neoformans*, mitochondrial transmission is influenced by mating-type or sexual-development regulators (Fedler et al., 2009; Yan et al., 2007; Sun et al., 2020). By contrast, in filamentous ascomycetes such as *Aspergillus flavus*, *Aspergillus nidulans*, or *N. crassa*, mitochondrial inheritance is more commonly described as maternal, meaning biased toward the parent that forms the female reproductive structure or sexual fruiting body rather than directly determined by mating-type identity (Coenen et al., 1996; Horn et al., 2016; Mitchell & Mitchell, 1952; Mannella et al., 1979). Mucorales are therefore particularly informative because they have mating types rather than morphologically defined male and female gametes (Idnurm et al., 2008), arguing that mitochondrial UPI in this group is not a passive consequence of unequal gamete size or anisogamy. Instead, it points to regulated sorting or elimination of one mitochondrial lineage during sexual development. By restoring homoplasmy after cytoplasmic mixing, mitochondrial UPI may reduce organelle conflict and help maintain cytonuclear compatibility in the recombinant progeny.

The evolutionary significance of this dual restriction is substantial. Biparental mitochondrial inheritance would generate heteroplasmy, resulting in intracellular competition between mitochondrial genotypes and potential conflicts between nuclear and mitochondrial genomes. By coupling a single meiotically reshuffled nuclear lineage to a single mitochondrial lineage, *M. circinelloides* may preserve cytonuclear compatibility while allowing recombination and chromosome assortment to occur without transmitting the full diversity of meiotic products. In this sense, sexual reproduction in Mucorales is not simply a generator of diversity. It is a highly filtered process in which recombination reshuffles nuclear chromosomes, but both nuclear and mitochondrial transmission are narrowed before viable progeny emerge.

## Materials and methods

### Fungal strains and culture conditions

The wildtype *Mucor circinelloides* phylogenetic species 15 (PS15) strains CBS394.68 and CBS172.27, corresponding to the (−) and (+) mating types, respectively, were used as parental strains throughout this study. Unless otherwise indicated (e.g., mating assays), strains were cultured on yeast-peptone-dextrose (YPD) medium at room temperature. Asexual spores were freshly harvested from cultures grown for 4–6 days.

### Mating assays, zygospore isolation, and progeny germination

Sexual crosses were performed as previously described (Pérez-Arques et al., 2025). Briefly, spores from the PS15 (−) and PS15 (+) parental strains were inoculated at four positions on Whey Agar (WA) plates and allowed to grow toward the opposite mating type at room temperature in darkness. After 7 days, mature zygospores formed at the interaction zone between the two mating types and were collected from the agar surface using sterile tweezers and suspended in distilled water.

Zygospore suspensions were first passed through a 100-µm cell strainer to remove larger hyphal debris. The filtrate was then passed through a 40-µm cell strainer, and the particles between approximately 40 and 100 µm were recovered to enrich for mature zygospores. Individual zygospores were manually isolated and transferred onto 1% agarose medium adjusted to pH 4.0. Plates were incubated at room temperature for 2–8 weeks and examined daily to allow completion of dormancy and zygospore germination. Zygospore germination was infrequent (1-2%).

Following germination, germsporangiospores were recovered in sterile water using a wetted micropipette tip. Spore suspensions were plated onto YPD medium adjusted to pH 3.2 to obtain isolated colonies derived from single germsporangiospores. Individual colonies were then selected for subsequent genetic and genomic analyses. In zygospores producing branched germsporangiophores, germsporangiospores from spatially separated germsporangia were collected independently and plated separately to test whether distinct branches contained genetically distinct meiotic products.

Mating types of the recovered progeny were determined by backcrossing individual isolates with the wildtype PS15 (−) and PS15 (+) parental tester strains on Whey Agar medium. Mating-type identity was assigned according to compatibility with the opposite mating-type tester, as determined by the formation of sexual reproductive structures at the interaction zone.

### Genomic DNA isolation and sequencing

Ultra-high-molecular-weight genomic DNA was isolated from PS15 (−) and PS15 (+) parental strains spores pregerminated for 6 h using a phenol/chloroform-based extraction method. Sequencing libraries were independently prepared without barcoding using the Oxford Nanopore Technologies Ultra-Long DNA Sequencing Kit V14 (SQK-ULK114) and sequenced on FLO-MIN114 flow cells using a MinION Mk1C device. Raw sequencing data were subsequently basecalled using Dorado v.1.0.1 in super-accuracy mode with the ‘dna_r10.4.1_e8.2_400bps_sup@v5.2.0’ model. For chromosome-level scaffolding, spores from the PS15 (−) and PS15 (+) parental strains were submitted to Phase Genomics (Seattle, WA, USA) for Hi-C library preparation and sequencing using the Proximo Hi-C Kit for fungi. Whole-cell spores were processed according to the manufacturer’s protocol, including crosslinking, cell lysis, chromatin digestion with DpnII, HinfI, DdeI, and MseI, proximity ligation, crosslink reversal, streptavidin enrichment, and preparation of dual-indexed Illumina sequencing libraries. Hi-C libraries were sequenced by Phase Genomics on an Illumina NovaSeq X Plus platform using 150-bp paired-end reads.

For whole-genome sequencing of meiotic progeny, genomic DNA was isolated from overnight YPD broth cultures using the Norgen Biotek Yeast/Fungi Genomic DNA Isolation Kit. Sequencing libraries were prepared using the Roche KAPA HyperPrep Kit according to the manufacturer’s instructions and sequenced as 150-bp paired-end reads. Because progeny sequencing was performed over several years, libraries were sequenced on either Illumina NovaSeq 6000 or NovaSeq X Plus platforms, depending on instrument availability at the time of sequencing.

### Genome assembly, polishing, and chromosome-level scaffolding

De novo genome assemblies were generated independently for PS15 (−) and PS15 (+) using ONT ultra-long sequencing reads. Nanopore reads were corrected and assembled using Canu v2.2, specifying a genome size of 38 Mb (genomeSize=38m) and a minimum read length of 5 kb (minReadLength=5000).

The resulting assemblies were polished through three successive rounds of Racon v1.5.0 using ONT reads. Before each polishing round, a genome-specific repetitive 15-mer database was generated using Meryl v1.4.1 (k=15), and repetitive k-mers were identified using a distinct-frequency threshold of 0.9998. Nanopore reads were subsequently aligned to the current assembly using Winnowmap v2.03 with the map-ont preset and the repetitive k-mer database supplied through the-W parameter. The resulting alignments were used by Racon to generate a corrected assembly, which served as the reference for read alignment and polishing during the following iteration.

Following long-read polishing, assemblies were further corrected using paired-end Illumina reads. Short reads were aligned to each assembly with BWA-MEM v0.7.17 using the-a option to retain alternative alignments. Alignments corresponding to the first and second reads of each pair were separated using SAMtools v1.10 and provided to Polypolish v0.6.0 for reference-guided correction. A subsequent round of short-read polishing was performed using Pypolca v.0.3.1 with paired-end Illumina reads and the --careful option.

For chromosome-level scaffolding, Hi-C paired-end reads were aligned to the corresponding polished parental assembly using BWA-MEM v.0.7.17 with the-5PM-T0 options. Alignments were converted to BAM format and coordinate-sorted using SAMtools v1.10. PCR duplicates were removed using Picard v.3.0.0 MarkDuplicates, and unmapped, secondary, and supplementary alignments were excluded using SAMtools (-F 2308). Alignments carrying alternative-hit (XA:Z:) tags were additionally discarded. The resulting alignments were sorted by read name and supplied to YaHS v1.2.2. for Hi-C-guided scaffolding. Scaffolding was performed using the restriction enzyme recognition sequences GATC, GANTC, CTNAG, and TTAA, and the canonical telomeric repeat motif TTAGGG. Scaffolding parameters were explored across multiple runs and the final assemblies were generated using ten scaffolding rounds (-R 10) and default resolution settings. The resulting chromosome-level assemblies were assessed for contiguity, correspondence with Hi-C linkage information, and the presence of telomeric repeats at chromosome termini.

Assembly contiguity and completeness were evaluated using QUAST v.5.2.0 and BUSCO v.5.7.1, respectively, including assessment of conserved single-copy eukaryotic orthologs using the eukaryota_odb10 dataset.

### Repetitive sequence and gene annotation

Repetitive sequences were characterized independently in both parental genome assemblies using complementary de novo and iterative repeat-classification approaches. Genome-specific repeat databases were constructed with BuildDatabase, and de novo repeat families were identified using RepeatModeler2 v2.0.6 through Dfam TE Tools v1.92, including structural identification of long terminal repeat retrotransposons with the-LTRStruct option. RepeatModeler2-generated libraries were separated into classified and unclassified repeat families, and unclassified elements were subjected to iterative classification. Classification was performed against fungal RepBase entries and previously classified repeat families, with newly identified elements incorporated into the classified repeat library during successive iterations.

Genome masking was subsequently performed through a serial RepeatMasker v4.1.9 workflow. Simple and low-complexity sequences were masked first, followed by known fungal repetitive elements, classified RepeatModeler2-derived families, and the remaining unclassified de novo repeat families. RepeatMasker was executed using the NCBI RMBlast search engine (-e ncbi), with low-complexity masking disabled in the subsequent repeat-library searches (-nolow). Repeat annotations generated during successive masking stages were combined into comprehensive repeat annotation files and converted to GFF3 format for downstream analyses.

As a complementary approach, repetitive and transposable elements were also identified using the Extensive de novo TE Annotator (EDTA) v2.2.2, with sensitive repeat detection and annotation enabled (--anno 1 --sensitive 1 --maxdiv 80 --species others). EDTA-derived annotations were used for chromosome-scale visualization.

Structural and functional gene annotation was performed independently for PS15 (−) and PS15 (+) using Funannotate v1.8.15. Genome assemblies were initially processed using funannotate clean, and repetitive regions were soft-masked using BEDTools v2.30.0 (maskfasta - soft) based on the corresponding repeat annotations.

Gene prediction was supported by paired-end RNA-sequencing data from previous studies (Pérez-Arques et al., 2025), which were processed independently against both parental assemblies using funannotate train. Gene models were subsequently predicted using funannotate predict and further refined using funannotate update. Functional annotation of predicted proteins was performed using eggNOG-mapper v2.1.12 with the eggNOG v5.0.2 database, with Gene Ontology evidence reporting enabled (--go_evidence all), and InterProScan v5.73-104.0, including InterPro domain assignments, Gene Ontology terms, and pathway annotations (--iprlookup, --goterms, and --pathways). The resulting functional annotations were incorporated into the gene annotation datasets using funannotate annotate.

### Chromosome characterization, synteny, and centromere identification

Chromosome-scale synteny between the PS15 (−) and PS15 (+) assemblies was assessed by aligning the corresponding chromosome sequences using Minimap2 with the assembly-to-assembly preset (-x asm5-N 0 --secondary=no). Chromosome alignments, gene annotations, repetitive sequences, and other genomic features were integrated for comparative visualization.

Ribosomal RNA genes were identified using Barrnap v0.9 and mapped onto the chromosome-level assemblies to characterize the genomic distribution of ribosomal DNA loci. Telomeric sequences were identified by searching for the canonical 5′-TTAGGG-3′ repeat and its reverse complement at chromosome termini.

Local GC-content variation was evaluated using BEDTools v2.30.0. GC content was calculated in nonoverlapping 200-bp genomic windows using bedtools makewindows and bedtools nuc. The deviation of each window from the genome-wide average GC content was calculated, and windows exhibiting absolute deviations greater than 15 percentage points were retained for visualization of regions with unusually high or low GC content.

Centromeric regions were identified by integrating previously generated H3K9me2 chromatin immunoprecipitation sequencing (ChIP-seq) profiles (Pérez-Arques et al., 2025) with sequence characteristics of Mucorales centromeres. ChIP-seq reads were quality-filtered using fastp v0.23.4 and aligned to the reference genome using BWA-MEM. H3K9me2-enriched regions were identified using MACS2 v2.2.9.1, and the H3K9me2 enrichment ratio relative to input DNA was quantified using bamCompare from the deepTools v3.5.6 suite. Centromere assignments were supported by the colocalization of H3K9me2-enriched heterochromatin, conserved centromere-associated DNA motifs, adjacent AT-rich regions, and clusters of Grem-LINE1 retrotransposons. The distribution of centromeres, rDNA loci, telomeric repeats, and repetitive elements was visualized across both parental chromosome assemblies using Circos v0.69-9.

### Genetic variation and meiotic recombination analyses

Illumina whole-genome sequencing reads from the parental strains and meiotic progeny were processed using fastp v0.23.4 to remove adapter sequences and low-quality reads, specifying a minimum average read quality of 20 and a minimum retained read length of 148 bp (-- average_qual 20 --length_required 148 --length_limit 0 --trim_poly_g --dup_calc_accuracy 1).

Processed reads were aligned to the chromosome-level PS15 (−) reference assembly using BWA-MEM with the-M option to maintain consistent genomic coordinates across parental and progeny datasets. Resulting alignments were coordinate-sorted, duplicate-marked, and assigned appropriate read-group information using GATK v4.4.0.0 and Picard v3.0.0 utilities. Reference genome indexes and sequence dictionaries were generated for downstream variant calling.

Single-nucleotide variants (SNVs) distinguishing the PS15 (−) and PS15 (+) parental strains were identified using GATK v4.4.0.0 HaplotypeCaller with the ploidy set to one (--sample-ploidy 1). Mixed and symbolic variants were excluded using SelectVariants. Low-confidence variants were identified using VariantFiltration with the following thresholds: quality by depth (QD) below 20, variant quality (QUAL) below 30, strand odds ratio (SOR) above 3, Fisher strand bias (FS) above 60, and mapping quality (MQ) below 40. Variants failing quality filters and variants shared between the parental strains were excluded.

Variants overlapping annotated repetitive regions were removed using BEDTools v2.30.0 (intersect-v), generating a high-confidence set of parental markers in nonrepetitive genomic regions. A total of 18,587 informative SNPs distinguishing the two parental genomes were retained for meiotic recombination and inheritance analyses.

To determine parental allele inheritance, sequencing reads from individual progeny were evaluated at informative SNV positions. Allele counts were extracted from aligned reads using GATK v4.4.0.0 GetPileupSummaries, and parental origin was assigned when at least 95% of reads supported either the PS15 (−) reference allele or the PS15 (+) alternate allele. Genomic positions with insufficient support for either parental allele were excluded from unambiguous inheritance assignments.

The distribution of inherited parental alleles was examined along each chromosome to identify meiotic crossover events and patterns of independent chromosome assortment. Consecutive informative markers originating from the same parent were grouped into haplotype blocks, with transitions between parental haplotypes interpreted as candidate meiotic recombination breakpoints. Genotypes of germsporangiospores derived from the same zygospore were compared across all informative markers to determine whether multiple meiotic products contributed to the viable sexual progeny.

### Mitochondrial genome assembly and inheritance analyses

Putative mitochondrial contigs were identified in the initial Canu assemblies based on their size and predicted circularity (suggestCircular=yes) and excluded from subsequent nuclear chromosome scaffolding. Circularity was assessed by inspecting terminal sequence overlaps, and redundant sequences were removed to obtain unit-length circular assemblies. In one assembly, a concatemeric contig containing approximately three tandem copies of the mitochondrial genome was resolved into a single-copy representation. Mitochondrial contigs were independently polished using parental Illumina short reads with Polypolish v0.6.0 and Pypolca v0.3.1.

Mitochondrial genomes were annotated using MFannot v1.37 with default parameters and the mold mitochondrial genetic code (translation table 4). MFannot annotations were converted into GFF3 format using AGAT, and mitochondrial genome organization, including gene distribution and repetitive elements, was visualized using Circos.

To identify parental mitochondrial polymorphisms, Illumina sequencing reads from PS15 (+) were mapped against the PS15 (−) mitochondrial reference genome, and variants were identified using GATK v4.4.0.0 as previously described. Variants were filtered using the following thresholds: QD < 20, QUAL < 30, SOR > 2, FS > 1, and MQ < 40. Only SNVs passing all quality filters were retained, yielding a total of 20 high-confidence, informative mitochondrial polymorphisms. Progeny sequencing reads were mapped against the same mitochondrial reference, and allele counts at informative SNV positions were determined using GATK v4.4.0.0 GetPileupSummaries to assess mitochondrial inheritance. Mitochondrial inheritance was assessed using the allele frequencies of informative SNVs to distinguish between uniparental and biparental transmission and identify the contributing mating-type parent. Uniparental inheritance was assigned when all informative SNVs showed greater than 95% allele support for the same parental genotype.

To establish a rapid, sequencing-independent method for assessing mitochondrial inheritance, a PCR-based restriction fragment length polymorphism (PCR-RFLP) assay was developed using a polymorphic restriction site distinguishing the parental mitochondrial genomes. HindIII was selected based on its widespread availability and the presence of a parental polymorphism within a readily amplifiable region, yielding diagnostic restriction fragments that could be clearly resolved by agarose gel electrophoresis. A 1,432-bp mitochondrial fragment was amplified using primers forward (5′-CACCTCTAGGAGATGCTCGAC-3′) and reverse (5′-ACCAATCCAAGGGATAGCCG-3′). PCR products were digested with HindIII and separated by agarose gel electrophoresis. The PS15 (−) mitochondrial genotype contained the HindIII recognition site, yielding fragments of approximately 1.02 and 0.41 kb, whereas the PS15 (+) genotype lacked this site and retained the undigested 1.43-kb amplicon. The assay was tested on a subset of progeny, with mitochondrial genotypes assigned according to their restriction profiles relative to the parental controls.

## Data availability

Previously published bulk progeny whole-genome sequencing, RNA-seq, and ChIP-seq datasets are available under BioProject PRJNA1170303 (Pérez-Arques et al., 2025). Newly generated sequencing data, including Oxford Nanopore, Hi-C, and individual progeny whole-genome sequencing, as well as the chromosome-level genome assemblies and annotations generated in this study, will be deposited and made publicly available upon peer-reviewed manuscript publication.

## Acknowledgements

We thank our lab manager, Anna Floyd Averette, for her support. We thank Dr. Devjanee Swain Lenz and Duke University’s Sequencing and Genomic Technologies Core Facility for their assistance with sequencing, as well as Thomas Milledge and the Duke Computer Cluster team for providing computational resources and support.

This study was supported by the National Institutes of Health/National Institute of Allergy and Infectious Diseases (NIH/NIAID) grants R01 AI050113-20, R01 AI039115-28, R01 AI170543-05, and R01 AI172541-04 awarded to J.H. C.P.-A. is supported by the Beatriz Galindo 2024 program (BG24/00103), funded by the Spanish Ministry of Science, Innovation and Universities and co-funded by the University of Murcia.

The funders had no role in study design, data collection and analysis, interpretation of results, or the decision to post this preprint. J.H. is a Fellow and Co-Director of the CIFAR program Fungal Kingdom: Threats & Opportunities.

